# Mammalian genomic regulatory regions predicted by utilizing human genomics, transcriptomics and epigenetics data

**DOI:** 10.1101/143990

**Authors:** Quan H. Nguyen, Ross L. Tellam, Marina Naval-Sanchez, Laercio R. Porto-Neto, William Barendse, Antonio Reverter, Benjamin Hayes, James Kijas, Brian P. Dalrymple

## Abstract

Genome sequences for hundreds of mammalian species are available, but an understanding of their genomic regulatory regions, which control gene expression, is only beginning. A comprehensive prediction of potential active regulatory regions is necessary to functionally study the roles of the majority of genomic variants in evolution, domestication, and animal production. We developed a computational method to predict regulatory DNA sequences (promoters, enhancers and transcription factor binding sites) in production animals (cows and pigs) and extended its broad applicability to other mammals. The method utilizes human regulatory features identified from thousands of tissues, cell lines, and experimental assays to find homologous regions that are conserved in sequences and genome organization and are enriched for regulatory elements in the genome sequences of other mammalian species. Importantly, we developed a filtering strategy, including a machine learning classification method, to utilize a very small number of species-specific experimental datasets available to select for the likely active regulatory regions. The method finds the optimal combination of sensitivity and accuracy to unbiasedly predict regulatory regions in mammalian species. Furthermore, we demonstrated the utility of the predicted regulatory datasets in cattle for prioritizing variants associated with multiple production and climate change adaptation traits, and identifying potential genome editing targets.

## Background

Predicting functional features of the genome beyond protein-coding regions has been the primary focus of the post-genome sequencing era [1, 2]. More than 90% of common genetic variants associated with phenotypic variation of complex traits are located in intergenic and intronic regions that regulate gene expression but do not change protein structure [3-5]. Moreover, SNPs associated with diseases such as autoimmune diseases, multiple sclerosis, Crohn’s disease, rheumatoid arthritis, and type one diabetes are strikingly enriched in promoters and enhancers [4, 6, 7]. Annotation of functional regions of the genome that harbour SNPs identified by genome-wide association studies (GWAS) to be significantly associated with variation in phenotype will contribute to the identification of functional SNPs and causative mutations, thereby suggesting genetic targets and markers for numerous applications in human health care and agricultural livestock production [8-11].

However, in mammalian species other than human and mouse, there is little data available at the genome level for discovery of regulatory elements. The recently established Functional Annotation of ANimal Genomes (FAANG) consortium has begun to address this deficiency in a coordinated fashion [12, 13]. It is expected that core assays identifying regulatory elements for key tissues in a number of production animals will be produced by the FAANG consortium and collaborators. However, the information generated in the foreseeable future for livestock is likely to remain far less comprehensive for coverage of tissues, sampling conditions and breadth of annotation of regulatory elements compared to human and mouse. The deficiency in the genome-wide prediction of regulatory elements is far greater for most other mammalian species. We have developed a computational method to utilize thousands of human regulatory datasets to predict regulatory elements in important mammalian species.

Transcriptional regulatory DNA elements (RDEs) are defined as genomic regions that are binding sites for one, or usually a combination of, transcription factors (TFs) and transcriptional coregulators [14-16]. Across distant species from *C. elegans* to *D. melanogaster* to humans, the architecture of gene regulatory networks, organization of chromatin topological domains, chromatin context at enhancer and promoter regions, and nucleosome positioning are remarkably conserved [17, 18]. For example, the majority of co-associations of transcription factors (i.e. combinations of different transcription factors binding to the same genomic region) at proximal transcription start site regions in human remain proximal in worm (80 %) and fly (100 %). Large-scale comparisons between humans and mouse *(M. musculus)* in the ENCODE project found a high level of conservation of binding motifs and activities, including: TF binding to different chromatin states (r = 0.9), proportion of enhancers in TF binding regions (r = 0.7), DNA methylation preferences within TF occupied regions (hypomethylated regions in both species), and TFs sharing conserved primary binding motif sequence (~94% of studied TFs) [19]. The human ENCODE, FANTOM, ROADMAP and related projects have generated large volumes of data relevant to the identification of promoters, enhancers and other RDEs [6, 20, 21]. However, these data have not been utilized for predicting regulatory genomic regions in other mammalian species – a strategy that can produce more comprehensive predictions than alternative options using a small set of experimental assays to identify a part of the regulatory repertory in the targeted species. We recognise that species specific regulatory elements may be underrepresented in this process. However, we note that the fundamental biology of, for example, that encompassing developmental programs, response to stimuli, reproduction, energy homeostasis, and many other systems show considerable conservation of components and processes across species [17, 19, 22].

In the current research, we developed the Human Projection of Regulatory Regions (HPRS) method to utilize results from thousands of biochemical assays in human samples to computationally predict equivalent information in other mammalian species. The method exploits the conservation of regulatory elements at the DNA sequence and genome organizational levels to map these elements to other mammalian species. It then uses species-specific data to filter these mapped sequences, which are enriched for regulatory sequence features, to predict a set of high confidence regulatory regions. We selected cattle as the target species to build the HPRS pipeline and then used the pig as a test species to validate the pipeline. The two species are important agricultural ruminant and non-ruminant species, respectively, with genomes sequenced but with little information available about genomic regulatory regions [12]. We also applied the method to the genomes of eight additional mammals. We demonstrated that the predicted regulatory dataset produced by the HPRS pipeline is useful for selecting more likely functional SNPs before (e.g. for SNP chip design) and after (e.g. for prioritising significant SNPs) GWAS analysis, genomic prediction models, and the understanding of biological mechanisms underlying non-coding genomic variant effects to potentially identify regulatory targets for genome editing.

## Results and Discussion

### A pipeline for the projection of human genomic features to other mammals

The four key elements of the HPRS pipeline (Fig. 1) include: (1) selection of suitable regulatory datatypes (biochemical assays) and tissues in humans; (2) mapping the selected features to the target species by utilizing conservation of genome organization and sequence identity to maximize coverage without compromising specificity; (3) first round filtering of the mapped regions to retain high-confidence mapped features, which had strict one to one forward and reciprocal mapping and where human features have multiple mappings to the target genome keeping only those with high sequence identity, and; (4) second round filtering by applying a pipeline to utilize available (often limited in scale and coverage) species-specific data to prioritize regions likely to be functional in the target species.

**Fig. 1.**
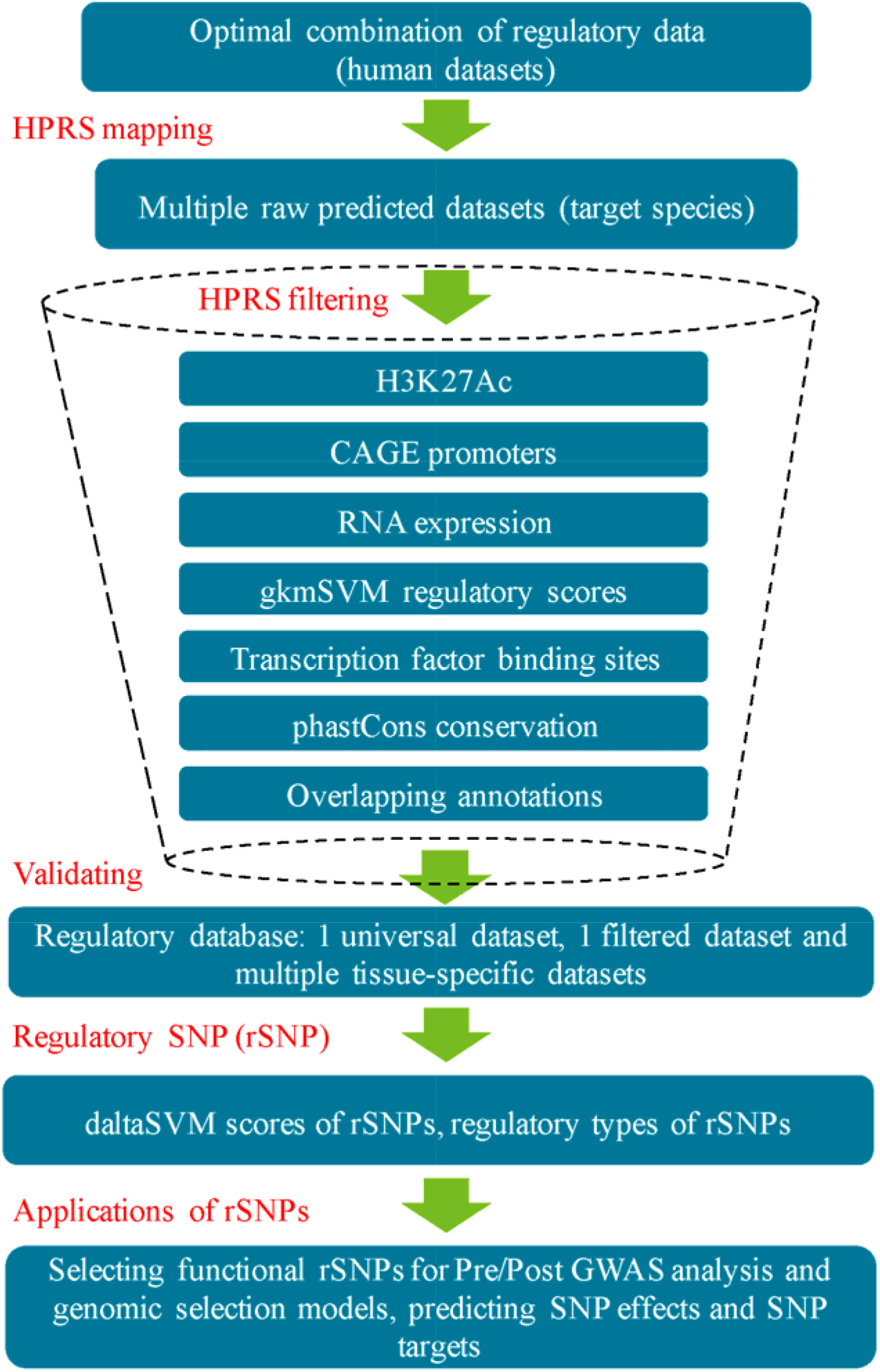
A streamlined workflow for the prediction of regulatory regions. Four key steps include mapping human regulatory regions to a target genome (creating a universal dataset), filtering the mapped regions by seven epigenomic, transcriptomic and genomic criteria to keep only regions with potential regulatory functions, validating the predicted regions by comparing with known reference dataset, and translating the findings to potential applications in genomic technology.

### Optimizing parameters for mapping sequence features across genomes

To identify regions that were likely to be orthologous between genomes we deployed the liftOver tool [23] and the precomputed alignment files available from the UCSC to map regulatory regions in the human genome to the cattle genome based on sequence similarity and genome location. First, we optimized the minMatch mapping threshold of the liftOver tool, which is the minimum proportion of bases to the total length of a region mappable to contiguous aligned segments in the target genome. The minMatch parameter was thoroughly tested with a range from high stringency 0.95 down to 0.1 (Fig. 2). The minMatch parameter values were assessed using seven diverse datasets (Fig. 2, Table 1).

**Fig. 2.**
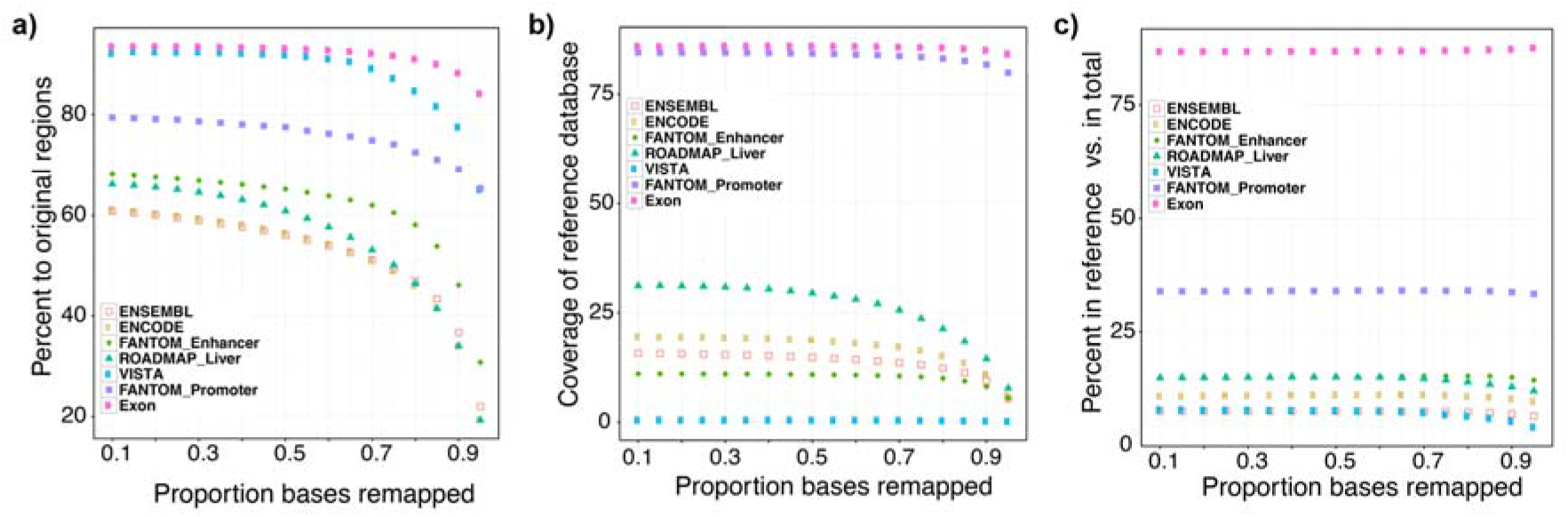
Optimization of mapping parameters using seven input databases. The input databases included five human enhancer databases (ENSEMBL, ENCODE, ROADMAP liver tissue, Vista, and FANTOM enhancers), one human promoter database (FANTOM promoters) and one annotated human exon database (UCSC hg19) [6, 18, 21, 56, 57]. The numbers of regions from each dataset used to optimise parameters is shown in Table S4. We used the UCSC pair-wise whole genome alignment chain files between the human genome (hg19) and the bovine genome (UMD3.1) and performed mapping from the human genome to the bovine genome (minMatch 0.1 to 0.95 as shown in the x-axis) and then reciprocal mapping from the bovine genome back to the human genome [52, 58-60]. a) recovered rate, defined as the percentage of the number of mapped regions with exact reciprocal mapping to the total number of original regions in humans. b) confirmation rate, defined as the percentage of reference regions covered by predicted regions to the total number in reference regions (Villar reference enhancers, Villar reference promoters, and cattle GENCODE genes V19). c) specificity, defined as the percentage of matched reference (true positive for the reference dataset) compared to the total number of predicted regions.

**Table 1.** Summary information for the optimised set of human regulatory datasets used for HPRS mapping^1^.

| Dataset | Tissues/cell lines | Total number of regions | Region types | Mean length (bp) |
| --- | --- | --- | --- | --- |
| ENCODE Distal TFs [19, 26] | 0/91 | 1,122,364 | Binding sites for 163 TFs | 151.2 |
| ENCODE Proximal TFs [19, 26] | 0/91 | 384,343 | Binding sites for 163 TFs | 151.4 |
| ROADMAP<sup>2</sup> [21] | 24 primary cells (e.g. blood cells, immune cells, and breast myoepithelial cells), 14 primary culture (e.g. skin, muscle satellite, neurosphere, bone marrow), and 50 primary tissues (e.g. thymus, spleen, lung, fetal stomach) | 9,102,278 | Enhancers | 970.8 |
| FANTOM Enhancers [6, 25] | 135/673 | 43,011 | Enhancers | 289 |
| FANTOM Promoters [20, 25] | 152/823 | 201,802 | Promoters | 21.5 |

<sup>1</sup>Information on data types and models is described in Table S3.
<sup>2</sup>See Table S5 for sample source details

The percentage of regions mappable to the target genome was compared to the total number of elements in the human regulatory databases (Fig. 2a). For cattle, mappable regions were defined as: 1) a small sequence segment (SSS) that can be mapped from the human to the bovine genome; 2) the resulting SSS can be mapped back (reciprocally mapped) from the bovine to the human genome; and 3) the boundaries of the reciprocally mapped SSS were within 25 bp of the boundaries of the original SSS in the human genome.

In all five enhancer datasets tested as shown in the Fig. 2a, the ratio of mapped regions increased steadily when the minMatch parameter was reduced from 0.95 to 0.55, with a much slower increase when the minMatch was reduced from 0.55 to 0.10 (Fig. 2a). The accuracy of the sequence projection was assessed as the percent of mapped regions that overlapped with a feature present in a reference cattle liver enhancer dataset, identified experimentally by histone 3 lysine 27 acetylation (H3K27Ac - a marker for active enhancers) and histone 3 lysine 4 trimethylation (H3K4me3 – a marker for active promoters near transcription start sites) assays (hereafter referred to as the Villar reference datasets) [22] (Fig. 2b). The coverage of the relevant reference datasets (Villar reference promoters, Villar reference enhancers and UCSC exons) also increased when the minMatch was reduced for some, but not all databases (Fig. 2b). Importantly, the reduction in mapping threshold did not lead to a loss of specificity, which is defined as the percentage of predicted enhancers that matched Villar reference enhancers (true positive for the reference dataset) compared to the total number of enhancers predicted using the particular input dataset (Fig. 2c). The combined results shown in Fig. 2a and Fig. 2b suggest that reducing minMatch to lower than 0.55 still increases (at a slower rate) the number of mapped regions (for the ROADMAP, ENSEMBL, FANTOM and ENCODE datasets – Fig. 2b) and increases the chance of detecting more reference enhancers (for the ROADMAP, ENSEMBL and ENCODE datasets (Fig. 2a). No significant difference was observed when lowering the minMatch from 0.2 to 0.1, but a slight gain in the percent of mappable regions was obtained when decreasing the minMatch from 0.3 to 0.2. Therefore, the parameter testing indicated that the optimal minMatch threshold was 0.2.

We also developed the method to detect regions possibly from gene duplication events (Supplementary Methods). To identify regions possibly resulted from duplication events (Fig. S1a), the HPRS mapping pipeline pooled unmapped regions in the human datasets (with minMatch=0.2) and mapped regions with no exact reciprocal matches for a second round mapping with different parameters (allowing multiple mappings and keeping only results with similarity higher than 80%) to rescue regions with multiple map targets.

### Optimised use of human regulatory datasets

Regulatory regions can be active or quiescent, depending on the cell type and the biological state, and therefore prediction using a single tissue/cell line, or a single assay type, is unlikely to produce a high coverage of all possible regulatory sequences of a species [24]. Therefore, we investigated the effect of using different databases on the predictive capacity of HPRS. First, we compared the mapping coverage of enhancers from 42 human ROADMAP datasets [21] to the reference liver enhancer datasets, which were experimentally identified (by H3K27Ac assay for liver tissues) for 10 mammalian species reported in Villar et al. [22] (Fig. 3a, 3b and Tables 1 and 5). Figure 3a shows the percentage of Villar reference enhancers (e.g. enhancers detected in liver tissues in cat) that overlap with HPRS predicted regions by mapping each of the original 42 human tissues to the targeted species (e.g. to the cat genome). Figure 3b shows the percentage overlapping with the results from using the combined 42 tissues. Comparing results from each tissue, or from combined tissues in each species, enabled assessment of variation due to evolutionary distance or tissue specificity. Second, we evaluated the predictions from human to bovine based on different datatypes, including: promoter databases (FANTOM), enhancer databases (FANTOM and ROADMAP), and transcription factor binding site databases (ENCODE proximal and distal TFs) (Fig. 3c, 3d). Each of the datatypes has unique sequence features that define different types of regulatory regions, for example those that are specific for promoters or enhancers. In general, species with closer evolutionary distance to humans had more HPRS predicted enhancers matching the relevant Villar reference liver datasets (Fig. 3a). For each tissue, the relative mapping rates were similar between species. Between different tissues across the 42 ROADMAP datasets, thymus enhancers having the lowest mapping rate and liver enhancers the highest mapping rate in most species (Fig. 3b). Notably, the tissue specificity effect, exemplified by the higher mapping rate for ROADMAP liver datasets to the relevant species Villar reference datasets than for other ROADMAP tissues (Fig. 3b), was reduced substantially if the two primates more evolutionarily related humans (macaque and marmoset) were removed from the comparison.

**Fig. 3.**
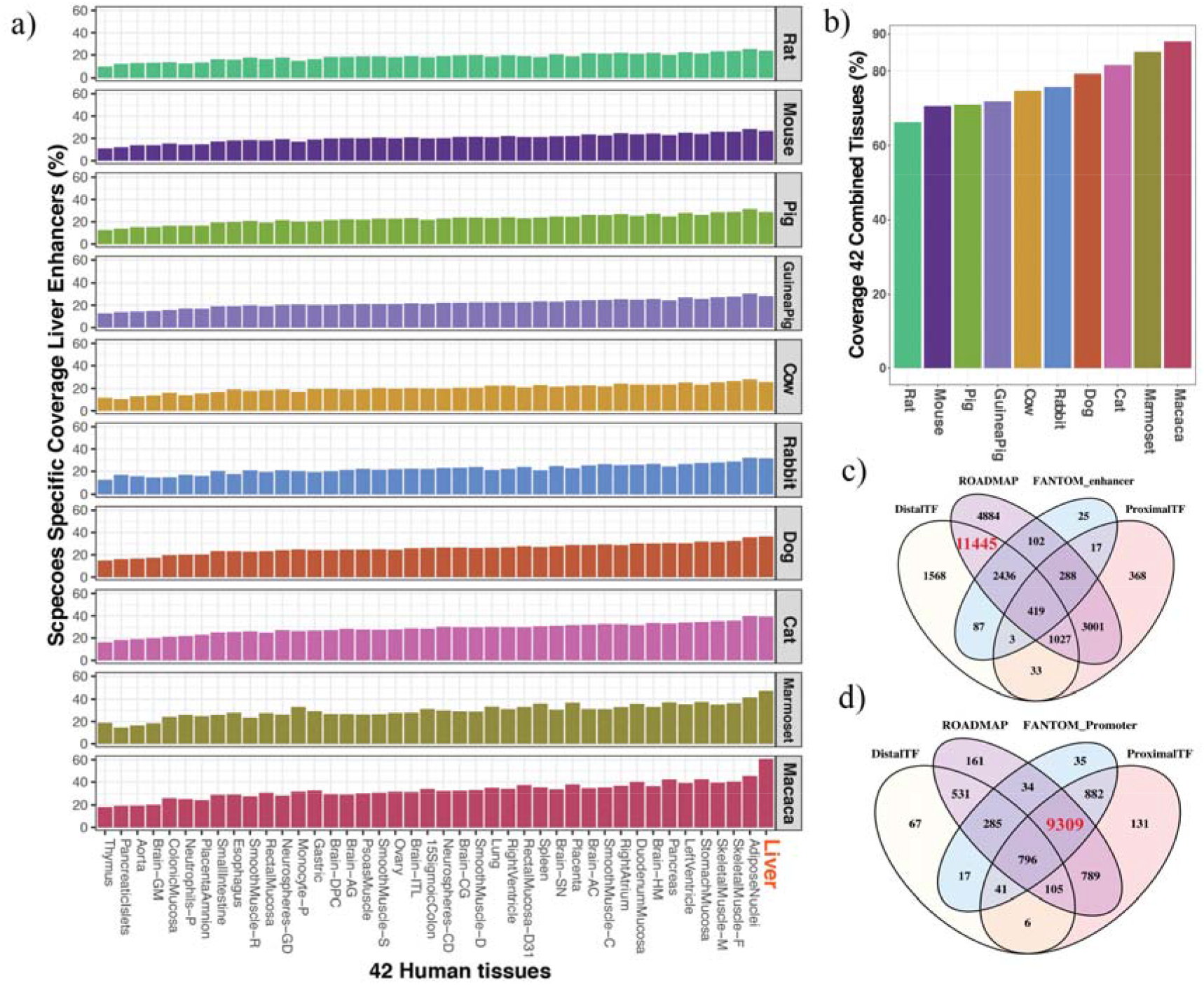
Effects of combining databases. **a)** Application of the HPRS mapping method to the genomes of ten mammalian species. The input for the HPRS mapping was one of the 42 ROADMAP human datasets [22] (38 adult tissues and four cell lines as shown in the x-axis) to 10 mammalian species. The Villar reference enhancer datasets determined by H3K27Ac and H3K4me3 assays for liver tissue in each species were used to estimate coverage of experimental enhancers by the predicted dataset (shown in y-axis as species-specific enhancer dataset). For each species, the coverage was the percent of the Villar reference enhancer dataset overlapped with the HRPS pre-filtered enhancers. **b)** The combination of all 40 tissues in each species was used. c) and d) show the optimal combination of five databases for enhancers and promoters respectively. The reference datasets include: ROADMAP enhancers (42 tissues); ENCODE distal TFs; ENCODE proximal TFs; FANTOM enhancers and FANTOM promoters. The numbers shown in the intersections are the number of common regulatory regions between the HPRS mapped regions and the Villar reference datasets.

**Table 2.** Summary of promoter predictions.

| Dataset | Total Regions in Cattle<sup>1</sup> | Overlap with Villar<sup>2</sup> dataset | Fold enrichment of Villar dataset | Number within 200 bp of TSSs<sup>3</sup> | Fold enrichment of TSSs |
| --- | --- | --- | --- | --- | --- |
| Total number CAGE regions | 154,377 (3.68 Mb, 0.138%) | 11,606 (84.1%) | 609 | 13,676 (51.0%) | 370 |
| Filtered set CAGE regions | 145,912 (3.46 Mb, 0.129%) | 11,203 (81.2%) | 629 | 13,011 (48.7%) | 377 |
| Total all regulatory regions (Universal Dataset) | 542,756 (937.39 Mb, 35.11%) | 13,329 (96.6%) | 3 | 20,759 (77.6%) | 2 |
| Filtered regulatory regions (Filtered Dataset) | 245,384 (356.1 Mb, 13.33%) | 13,104 (95.0%) | 7 | 17,715 (66.2%) | 5 |
| Villar reference promoters | 13,796 (32.90 Mb, 1.23%) | 13,796 (100%) | NA | 10,212 (38.2%) | 31 |
| ROADMAP promoters | 81,892 (135.6 Mb, 5.08%) | 12677 (91.9%) | 18 | 14,388 (53.8%) | 11 |

<sup>1</sup> The total number of regions with liftOver at minMatch 0.2 and exact reciprocal matches, combined with regions that had multiple matches (no one-to-one relationship) but had high conservation (80% identity). The original human regions are from the FANTOM promoter dataset. The percent was calculated for total genome size.
<sup>2</sup> Villar promoter dataset for cattle [22]
<sup>3</sup> Promoter count within 200 bp of the Ensembl annotated UMD3.1 TSSs Ensembl build 85 (total 26740).

**Table 3.** Summary of mapped and filtered regulatory sequences.

| Datasets | Number of mapped regions |  |  | Genome coverage (%) |  |  |
| --- | --- | --- | --- | --- | --- | --- |
|  | human | cow | pig | human | cow | pig |
| Total genome size (Mb) | NA<sup>1</sup> | NA | NA | 3,137.2 Mb<br>(100%) | 2,670.4 Mb<br>(100%) | 2,808.5 Mb<br>(100%) |
| ROADMAP enhancers (%<br>mapped to target species) | 9,102,278<br>(100%) | 5,917,129<br>(65%) | 5,620,417<br>(62%) | 8,836.6 Mb<sup>2</sup><br>(NA) | 6,142.4 Mb<sup>2</sup><br>(NA) | 5,809.5 Mb<sup>2</sup><br>(NA) |
| ROADMAP enhancers<br>(overlapping regions were<br>merged) | 494,583<br>(100%) | 371,295<br>(75%) | 361,682<br>(73%) | 1,123.2 Mb<br>(35.8%) | 885.6 Mb<br>(33.2%) | 826.2 Mb<br>(29.4%) |
| FANTOM CAGE enhancers | 43,011<br>(100%) | 34,303 (80%) | 27,558 (64%) | 12.4 Mb<br>(0.40%) | 12.2 Mb (4.6%) | 9.6 Mb (0.34%) |
| ENCODE distal TFs | 1,122,364<br>(100%) | 749,572<br>(67%) | 716,515<br>(64%) | 169.7 Mb<br>(5.4%) | 132.0 Mb<br>(4.9%) | 124.4 Mb (4.4%) |
| FANTOM CAGE promoter<br>peaks | 201,802<br>(100%) | 154,377<br>(76%) | 153,893<br>(76%) | 4.3 Mb<br>(0.14%) | 3.7 Mb (0.14%) | 3.7 Mb (0.13%) |
| ENCODE proximal TFs | 384,343<br>(100%) | 298,554<br>(78%) | 279,774<br>(73%) | 58.2 Mb<br>(1.9%) | 48.0 Mb<br>(1.8%) | 48.9 Mb (1.7%) |
| Merged ROADMAP,<br>ENCODE, and FANTOM<br>datasets (Universal Dataset)<sup>3</sup> | 760,702 | 542,756<br>(86.1% and<br>96.6%)<sup>4</sup> | 519,913<br>(89.2% and<br>97.1%)<sup>4</sup> | 1,165.7 Mb<br>(37.2%) | 919.5 Mb<br>(34.4%) | 857.8 Mb<br>(30.5%) |
| Filtered Dataset | NA | 245,384<br>(73.5% and<br>95.0%)<sup>4</sup> | 151,523<br>(69.8% and<br>95.6%)<sup>4</sup> | NA | 356.1 Mb<br>(13.3%) | 311.5 Mb<br>(11.1%) |

<sup>1</sup>NA, not applicable. The percent was not calculated for these three values because overlapping regions are present in the different enhancer datasets.
<sup>2</sup>The total size is bigger than the genome size because overlapping regions are included
<sup>3</sup>Total size of non-overlapping regions in the Universal Dataset (before filtering). The % overlapping Villar reference enhancers (the former) and promoters (the later) in the targeted species
<sup>4</sup>% overlap Villar reference liver enhancers and promoters in the filtered datasets

**Table 4.**
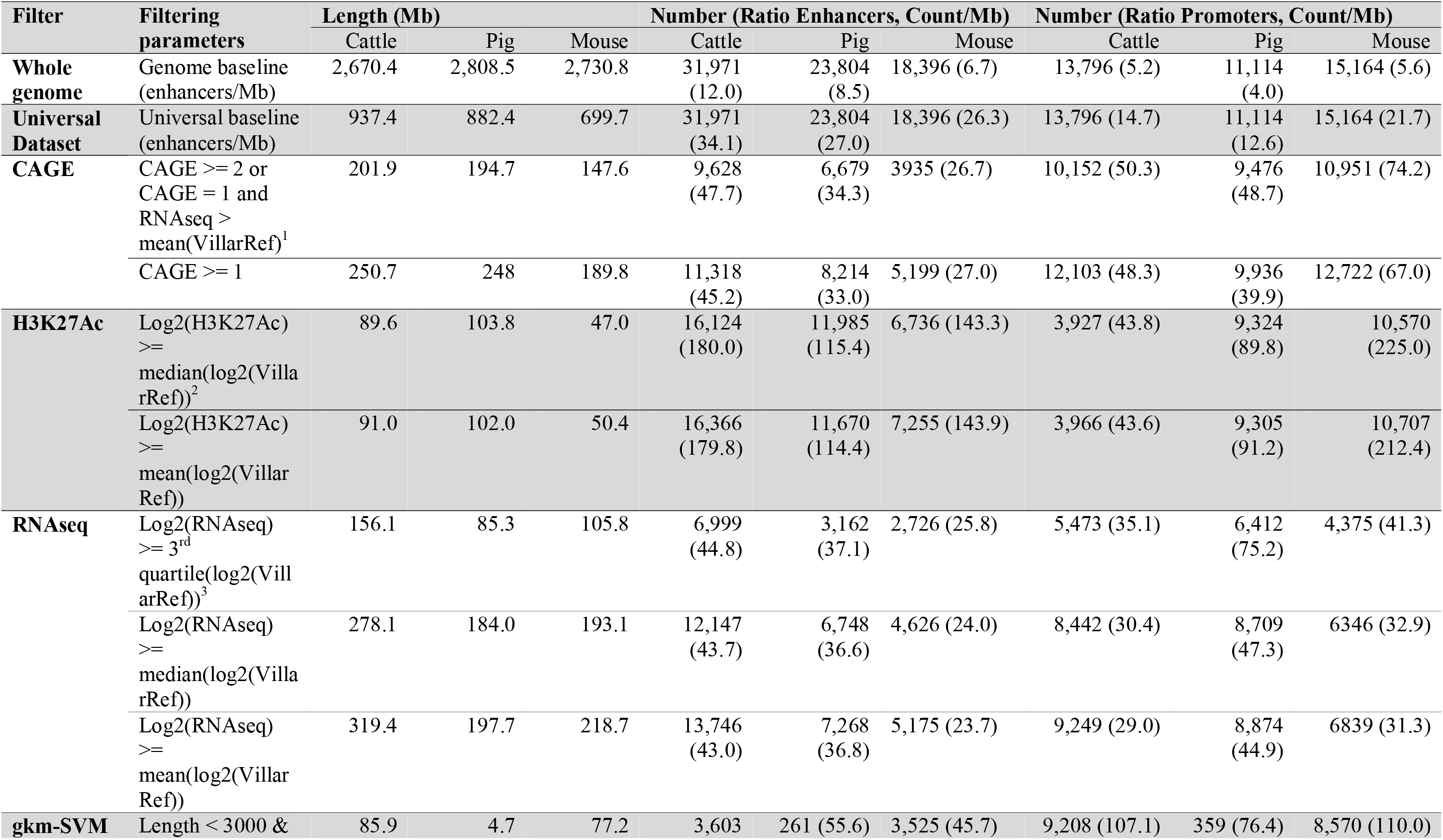
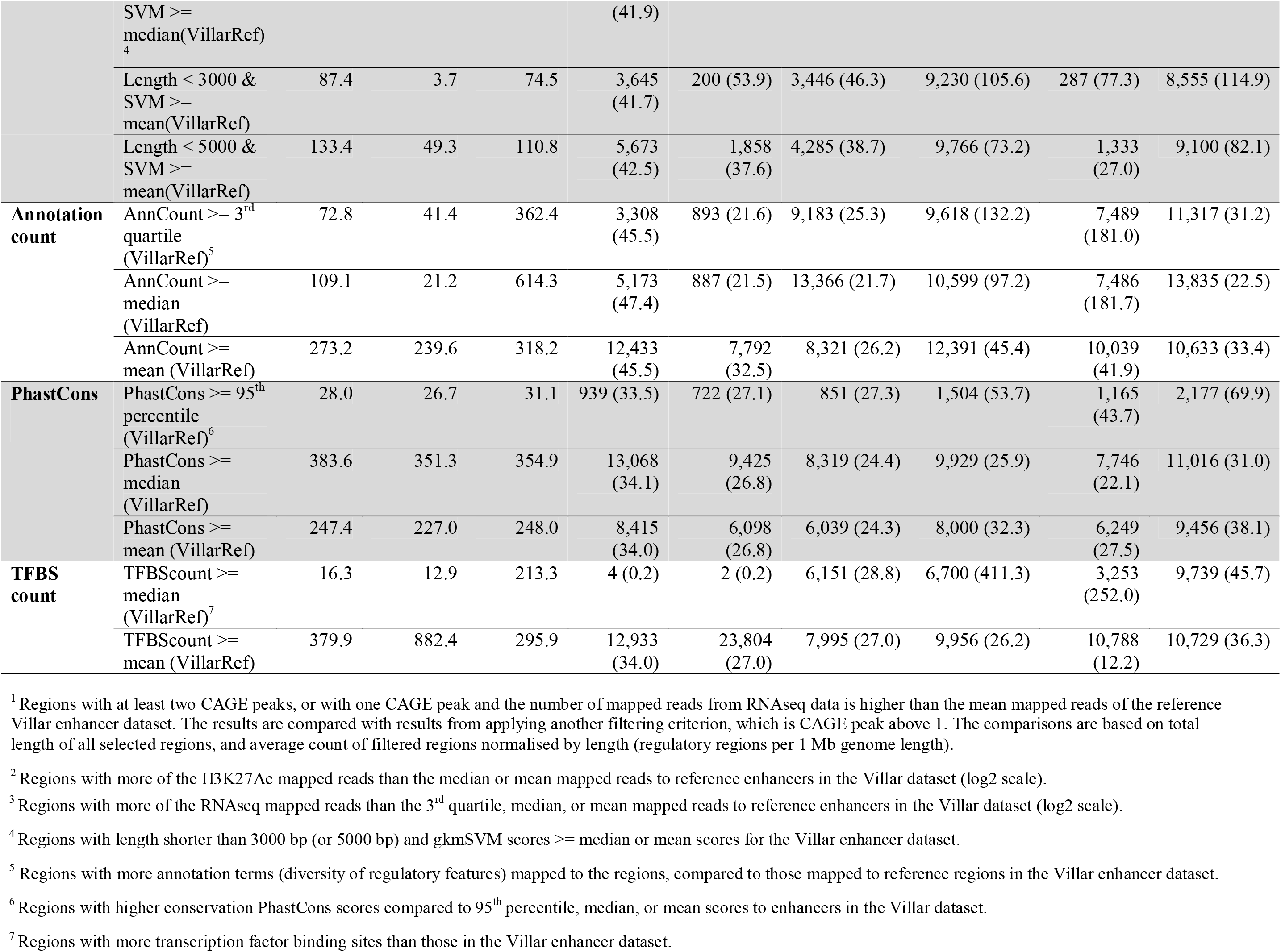
Filters with species-specific data for selecting regulatory regions (refer to the Supplementary Materials and Methods).

**Table 5.** HPRS predicted regulatory datasets for 10 species.

| Species | Number of regions | Total length (Mb) | Enhancer coverage<sup>1</sup> | Promoter coverage<sup>1</sup> |
| --- | --- | --- | --- | --- |
| **Unfiltered datasets** |  |  |  |  |
| Cattle (bTau6)<sup>2</sup> | 545,748 | 919.5 | 86.1% | 96.6% |
| Pig (susScr3) | 519,913 | 882.4 | 89.2% | 97.1% |
| Marmoset (CalJac3) | 642,144 | 1,106.4 | 93.1% | 98.4% |
| Rhesus Macaque (RheMac3) | 693,312 | 1,158.2 | 94.5% | 97.6% |
| Dog (CanFam3) | 570,317 | 877.5 | 89.4% | 97.6% |
| Cat (FelCat5) | 570,282 | 903.9 | 90.8% | 97.1% |
| Guinea pig (CavPor3) | 523,273 | 761.6 | 81.1% | 92.7% |
| Rabbit (OryCun2) | 531,109 | 819.4 | 86.8% | 96.8% |
| Mouse (Mm10) | 478,974 | 699.7 | 79.6% | 93.2% |
| Rat (Rn5) | 453,017 | 620.5 | 75.3% | 89.5% |
| **Filtered datasets<sup>3</sup>** |  |  |  |  |
| Cattle (bTau6) | 245,358 | 356.1 | 73.5% | 95.0% |
| Pig (susScr3) | 151,523 | 311.5 | 69.8% | 95.6% |
| Mouse (mm10) | 281,071 | 308.4 | 68.9% | 91.4% |

<sup>1</sup>Coverage of the relevant Villar reference datasets [22].
<sup>2</sup>Reference genomes are from UCSC (http://hgdownload.cse.ucsc.edu/downloads.html)
<sup>3</sup>The relevant Villar reference species enhancer datasets were added prior to filtering.

Since the coverage of the reference cattle liver enhancer dataset was not significantly higher with human liver enhancers, than with enhancers from many of the other human ROADMAP tissue enhancer datasets [21], we asked whether combining tissues would increase coverage. By combining the predictions from the 42 ROADMAP datasets, 2 to 4-fold higher coverage could be obtained than from one tissue alone (at least 60% total coverage) across a variety of species could be obtained, with coverage lowest for rat and highest for macaque (Fig. 3a, b). Furthermore, we found that separate databases constructed using different models and biochemical assays were complementary, and combining them significantly increased coverage compared with a single database alone (Fig. 3c, d). For example, prediction using the ENCODE distal TF dataset and the ROADMAP enhancer dataset covered the highest number of Villar cattle reference enhancers, while prediction using the FANTOM promoter and the ENCODE proximal TFBS databases covered more Villar cattle reference promoters, and each dataset could add a number of unique regulatory regions not found in other datasets (Fig. 3c, d). The combination of 88 ROADMAP datasets [21], the FANTOM enhancer and promoter datasets [25], and the ENCODE distal and proximal TF datasets [26] generated a maximum enhancer coverage of 95% (for macaque) and promoter coverage of 98% (for marmoset). Therefore, we selected an optimal combination of human input databases for the HPRS pipeline on the basis that they represent promoters, enhancers and TFBSs from a large combination of human tissues and primary cells and were generated by different methods (Table 1).

### Predicting promoters

One of the most comprehensive human promoter datasets is the FANTOM5 promoter atlas generated experimentally by CAGE data from almost one thousand tissues and cell lines [20]. CAGE is a sensitive methodology for the detection of transcription start sites (TSSs) and hence defines core promoter regions where there is binding of the transcriptional machinery [27]. Promoters generally have a high concentration of TFBSs, typically within 300 bp upstream and 100 bp downstream of the TSSs [20]. Promoter sequences are more evolutionarily conserved than enhancer sequences, and therefore a larger proportion can be mapped from human to other mammal genomes [22].

Of 201,802 CAGE transcription initiation peaks in the FANTOM5 human promoter atlas [20], 154,377 (76.5% of the total) were mappable to the bovine genome (Table 2). The HPRS using CAGE predicted new TSSs not present within the existing bovine genome annotation (Ensembl Build 85). Although a promoter dataset for cattle can be inferred by defining upstream sequences of genes with annotated TSSs, this indirect inference results in a small number of promoters. Approximately 26,740 cattle genes (coding, lncRNAs, miRNAs etc.) in the reference dataset used (Ensembl Build 85) have annotated TSSs. This dataset is far from comprehensive because the expected underrepresentation of non-coding genes and of alternative promoters (AP). The one gene-one promoter and one gene-one protein concepts are no longer appropriate to describe the diverse transcriptome [28]. AP are common and are functionally important. A number of APs were found associated with complex traits [29]. While 51% of the Ensembl cattle TSSs are covered by mapped human CAGE transcription initiation peaks (3.7 Mb), only 38.4% are covered by the experimentally defined promoters (32.9 Mb) in Villar et al. [22], suggesting that HPRS predictions based on human CAGE data could enrich promoter coverage in the cow by more than 12 times compared to the standard promoter assay (H3K4me3 ChIP-Seq) (Table 2). Active TSS regions from 88 human tissues in the ROADMAP were mapped to 81,892 putative promoters in cattle [21], with a total length of 135.6 Mb. Noticeably, the average number of Ensembl reference TSSs overlapped to every 1 Mb of predicted promoters based on the ROADMAP database was 37-fold lower than those based on the CAGE database (Table 2).

HPRS using the CAGE dataset can predict many TSSs at single-nucleotide resolution and can accurately predict transcriptional orientation. TSSs are presented in the Ensembl database as single nucleotide genomic positions. HPRS predicted promoters based on CAGE had exact overlap to the 7,191 Ensembl TSSs for cattle. While promoter prediction by using histone marks (such as those used by ROADMAP) cannot directly define transcriptional orientation, this information predicted by HPRS using human CAGE data is highly accurate [20]. Consistently, we found that of 13,676 genes that have TSSs within 500 bp of mapped CAGE peaks, 96.9% (13,257) genes had the same transcriptional orientation in the Ensembl annotation and predicted by human CAGE data. We therefore assigned promoter orientation using the predictions from the CAGE dataset.

### Mapping enhancer datasets

Prediction of enhancers is likely to be more challenging than predicting promoters because: 1) enhancers are less conserved in DNA sequence; 2) enhancer locations evolve faster [19, 22], and 3) enhancer effects are usually independent of the distance, orientation, and relative location (upstream or downstream) of gene targets [14]. To predict a broad set of sequences in a species that are active in one or more tissues or conditions, we expanded the human enhancer datasets to include: 88 tissues, primary cell lines and primary cell cultures generated by the ROADMAP project [21] (Table 1); all human active enhancers defined by CAGE data from hundreds of tissues and cell lines in the FANTOM project [6], and; all the Villar experimentally defined reference cattle liver enhancers [22] (Table 1). Cumulatively, the HPRS pipeline mapped over 9.1 million human enhancer sequences to over 5.9 million regions in the bovine genome, which were then merged into 542,756 non-overlapping regions (Table 3). The merged dataset (Universal Dataset) covered 86% (excluding merged regions resulting from the original Villar reference enhancers) of the Villar enhancer reference dataset (Table 3). The term “Universal” reflects the initial pooling of all relevant human regulatory datatypes and datasets to form a large collection of genomic regions to be mapped to the target species. Regulatory sequences are often active in certain conditions, and remain inactive in most other cases. Therefore, pooling active regulatory regions from a large number of datasets can likely cover most active and inactive regulatory sequences, thus enabling the prediction of a Universal Dataset.

The HPRS mapping of the enhancer datasets predicted a large set of homologous regions that are potentially regulatory regions in cattle (the Universal Dataset). We noted that alignability of DNA sequence does not automatically imply functionality [22], and therefore we applied a filtering pipeline to incorporate other types of cattle-specific data to prioritize regions more likely to have transcription regulation functions. The filtering pipeline used a combination of sequence features and epigenetics marks to enrich for likely regulatory enhancers and promoters, as discussed in the filtering section.

### Mapping transcription factor binding site datasets

To include potential regulatory regions beyond typical promoter and enhancer classifications, we performed HPRS mapping of human experimentally defined ENCODE TFBSs (ENCODE annotation version 2) to the bovine genome. The ENCODE TFBS database contains binding sites for 163 key TFs, some of which represent additional types of regulatory regions other than enhancers and promoters [30] (Table 1). The use of these TFBS datasets not only supported predictions from using the enhancer and promoter datasets, but more importantly added other regulatory categories into the combined prediction of regulatory regions. For example, the binding targets of the CCCTC-binding factor (CTCF) are likely insulator regions [31], while enhancer of zeste homolog 2 (EZH2) binding sites may mark polycomb repressor complex 2 (PRC2) regions [32]. These ENCODE TFBSs were identified as binding regions of TFs to nucleosome free regions (~151 bp per region), which are more biologically relevant than de novo scanning of genome sequence for TFBSs based on short position weight matrices (PWMs, typically 6-12 bp) because the later method only uses DNA sequence and does not take into account the biological chromatin context, which is essential for transcription factor binding [33, 34]. In total, from the ENCODE TFBS dataset [26, 34], 298,554 proximal TFBSs (total 47.97 Mb), and 749,572 distal TFBSs (total 132.04 Mb) were projected by HPRS onto the bovine genome. We also show that the HPRS prediction using ENCODE transcription factor datasets was supported by two other independent prediction approaches (Supplementary Methods).

### The filtering pipeline for a high-confidence regulatory region dataset

The predictions produced by HPRS were optimized so that they occupied a relatively small part of the whole genome, but can universally predict regulatory regions in different cell types and tissues. Applying HPRS for selected datasets (Fig. 3 and Table 1), we first produced a preliminary Universal Dataset then refined it to generate a Filtered Dataset (Table 3). To remove redundancies, overlapping mapped ROADMAP enhancers (initially mapped separately for each of the 88 ROADMAP datasets) were merged (Table 3). Similarly, all mapped regions for promoters, merged enhancers and TFBS with overlapping coordinates were merged into larger regions to form the final Universal Dataset (UD), containing 542,756 non-overlapping regions. These regions covered 937.4 Mb (35.1%) of the bovine genome. The high coverage (35.1%) of the UD was due to the large collection of human datasets used as inputs for mapping to bovine (37.2% of the human genome) so that the UD covered almost all possible promoters, enhancers and TFBS (Table 3). Importantly, the HPRS pipeline improves the specificity of the UD by applying a filtering step, which incorporates the power of cattle specific data to predict a small set of potential transcription regulatory genomic regions in the bovine genome (Fig. 4, Table 4).

**Figure 4.**
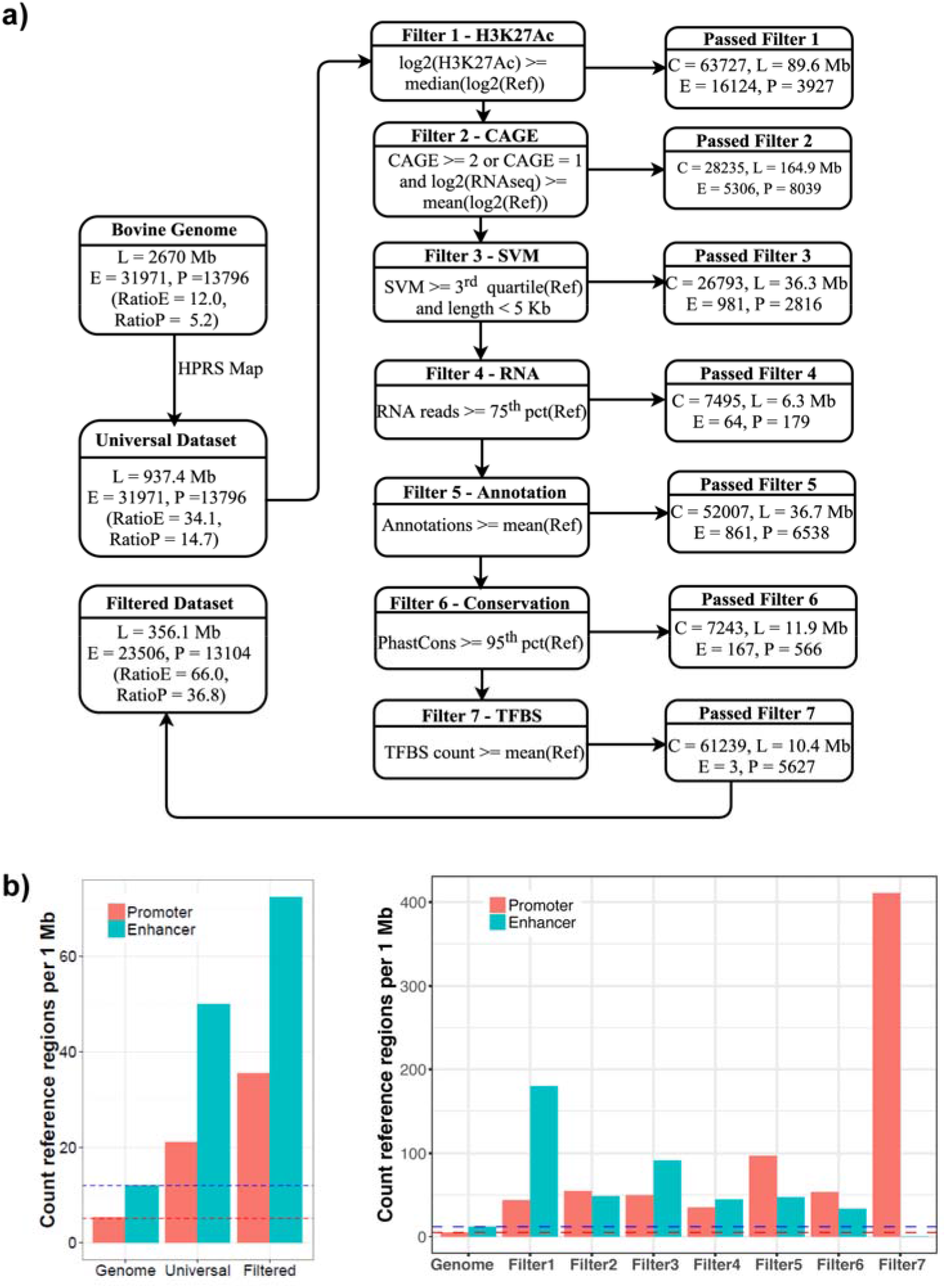
Enrichment of the enhancers and promoters by the filters in the HPRS filtering process. **a)** A pipeline to filter predicted regulatory regions from the Universal Dataset with 542,756 regions, covering 937.4 Mb of the genome (35.1%). The initial number of experimentally defined Villar reference datasets included 31,971 enhancers (E) and 12,257 promoters (P). The number of reference E and P, total number of predicted regulatory regions and total length (in Mb) for all promoters and enhancers passing each filtering layer are shown. The RatioE (total enhancers overlapping Villar reference enhancers/total length) and RatioP (total promoters overlapping Villar reference promoters/total length) were used as criteria to assess enrichment for each filter. The seven filters are described in the Table 4 and in the supplementary methods. **b)** Two bar graphs showing enrichment results (using the same starting set) of using each of the seven filtering steps in comparison with the baseline (whole genome) as shown in the dashed lines, and the Universal Dataset (mapped regions, not filtered). The y-axis shows the average number of reference promoters or enhancers in every 1MB of the genome. The density of regulatory regions predicted is an indicator of the prediction coverage and accuracy. The higher values indicate more experimentally validated enhancers and promoters are enriched after filtering, suggestive of a more efficient filter. Each filter was tested independently, using the same Universal Dataset as the input, to compare enrichment levels resulted from each of the seven filters.

The filtering pipeline reduced the UD to the much smaller Filtered Dataset (FD, the same as filtered UD) which covered a smaller part of the whole genome, but which still predicted most active enhancers and promoters (Table 4 and Figure 4). Detailed discussion on rationale for selecting each filter is in the Supplementary Materials and Methods. Briefly, the pipeline utilized both biological data in the target species (86 RNA-Seq datasets representing 79 cattle tissues [35], cattle H3K27Ac signal [22], and DNA sequence conservation scores) and computationally estimated criteria (gapped k-mers support vector machine (gkm-SVM) scores, number of overlapping annotations and number of CB-predicted TFBS) (Fig. 4a).

Before filtering, the Universal Dataset had approximately 2.84 times higher Ratio_E_ (Number of Villar reference enhancers by predicted regions divided by the length in Mb of predicted regions) and 2.82 higher Ratio_P_ (Similar to Ratio_E_, but for promoters) than the total genome baseline and each filtering step in the pipeline increased RatioE and RatioP compared to the baseline (Fig. 4b, Table 4). At the end of the pipeline, a set of high-confident regulatory regions (the FD), containing 245,384 sequences (with total length 356.1 Mb, equivalent to 13.3% of the whole genome) was obtained. The filtering reduced the number of regions by 2.2 times and the genome coverage by 2.6 times (Table 3, Fig. 4a), while still including most of the cattle liver reference enhancers and promoters (73.5% and 95.0% respectively) (Table 4, Fig. 3a). Importantly, the filtered dataset had a 5.5 and 7.1 times higher Ratio_E_ and Ratio_P_, respectively, than the genome baseline (Fig. 4). The size and coverage of the bovine genome (356.1 Mb, 13.3%) by HPRS predicted regulatory regions was comparable to the published figure for mouse, which is 12.6% of the mouse genome, as predicted by ENCODE DNAse I accessibility data and transcription factor ChIP-Seq (using antibodies for 37 TFs on 33 tissues/cell lines) and histone modification ChIP-Seq data [2]. Similarly, applying the HPRS pipeline to the mouse genome, without using mouse-specific datasets from ENCODE or other sources (except for the reference Villar dataset), predicted potential regulatory regions that occupy 11.3% of the whole mouse genome.

### Validating and extending the HPRS pipeline in nine other mammalian species

The performance of the HPRS pipeline was evaluated using the porcine (pig) genome (susScr3) [36]. HPRS had been developed based on the bovine genome, and the pig was then selected as a species for step-by-step comparison throughout the pipeline because of the availability of experimentally defined porcine promoter and enhancer reference datasets [22] and because the pig is an evolutionarily divergent non-ruminant production animal. We obtained similar results in pig compared to cattle on: numbers of putative regulatory regions, percent to total genome length, coverage of the reference datasets (Table 3 and Table 4). Importantly, we extended the application of the HPRS mapping data from human to 8 additional mammalian species, which had reference promoter and enhancer datasets from the Villar et al study. We generated HPRS mapped unfiltered UDs and observed consistently high coverage of the reference enhancer and promoter datasets and the coverages were comparable between all 10 mammalian species (Table 5). Thus, the pipeline appears to have general utility, not just for livestock species, but also for mammals in general.

### SNPs in regulatory regions are enriched for significant GWAS SNPs

Over 90% of significant GWAS SNPs lie outside gene-coding regions, and for those within the gene-body (from the start to the termination site of the complete transcript, including introns), over 92% are within intronic regions [3, 5]. To test the enrichment of potential causal SNPs within predicted regulatory regions in cattle, we explored the overlap between SNPs in regulatory regions and pleiotropic SNPs, which are SNPs significantly associated with multiple traits. The pleiotropic SNPs were identified by an independent GWAS study for 32 cattle feed intake, growth, body composition and reproduction traits [37]. The GWAS used 10,191 beef cattle, with data (including imputed data) for 729,068 SNPs (Fig. 5). We observed a substantial fold enrichment (~2-4 times) of SNPs with –log(*P*-value) from 3 to 20 in the Filtered Dataset compared to all other sets of commonly classifying SNPs in different genomic regions, including the set of SNPs 5 kb upstream of protein coding genes. We also observed higher counts (for 6 out of 10 traits) of associated SNPs within regulatory regions in a study on ten climatic adaptation traits in 2,112 Brahman beef cattle [38] (Fig. S1). Similarly we found enrichment of regulatory SNPs in a study of five major production and functional traits in 17,925 Holstein and Jersey dairy cattle (*p* < 0.05 for 3 out of 5 traits) [39] (Table S1). These observations are consistent with the pipeline identifying regulatory SNPs from millions of SNPs in the genome and suggest that the predicted regulatory database is useful for prioritizing SNPs likely to be contributing to phenotypic variation of complex traits.

**Figure 5.**
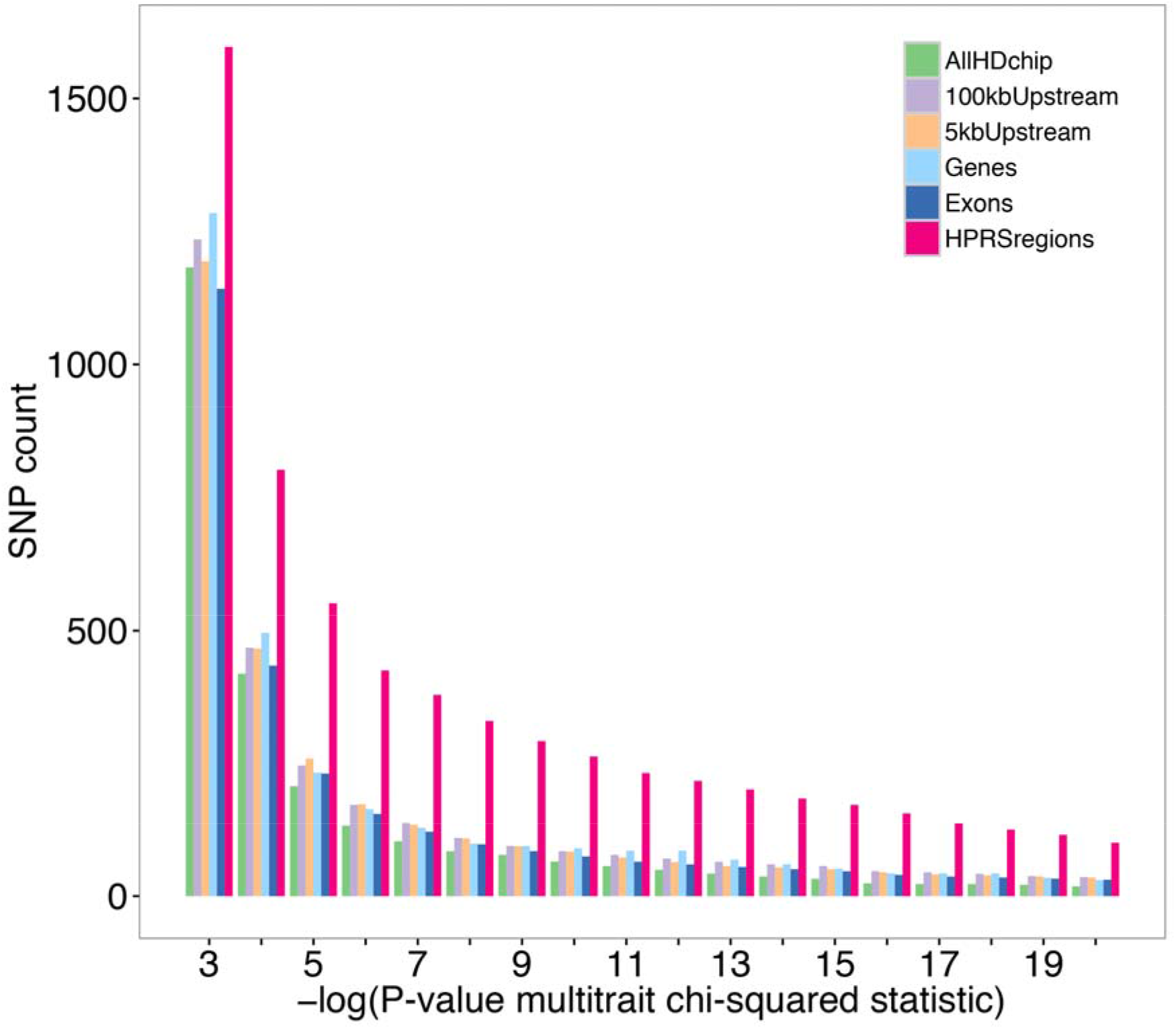
Enrichment of significant pleiotropic SNPs in regulatory genomic regions. Count of significant pleiotropic GWAS SNPs [37] (*P-values* are from multi-trait metaanalysis chi-squared test statistic for 32 traits) in a set of ~729,100 SNPs genotyped using the Illumina HD Bovine SNP chip or imputed from genotyping data from smaller Illumina Bovine SNP chips. Legend labels, from top to bottom: “AllHDchip”: 43,130 SNPs randomly selected (from all 692,529 SNPs on the HD chip, excluding those from chromosome X); “100kbUpstream “: 43,130 SNPs randomly selected (from 325,227 SNPs within 100 kb upstream regions of coding genes); “5kbUpstream”: all 30,384 SNPs within the 5kb upstream regions of coding genes (results scaled to 43k SNPs); “Genes”: 43,130 SNPs randomly selected (from 240,160 SNPs in coding genes); “Exons”: all 10,003 SNPs in exons of coding genes (results scaled to 43k SNPs); “HPRS regions”: 43,130 SNPs in regulatory regions.

### The regulatory region datasets can be used to guide identification of potential causative SNPs and their gene targets

As examples of the application of our resources to identify likely causative mutations from a large list of significantly associated SNPs, we applied the HPRS approach to analyse two well studied genetic variants in cattle, which were known to contribute to phenotypic variation, but their mechanisms of action were not known because they were located within non-coding regions.

The bovine Pleomorphic adenoma gene 1 (*PLAG1*) locus has been identified in the control of stature (weight and height) by several independent GWAS studies in cattle [40, 41]. The study by Karim et al. [40] fine-mapped 14 SNPs associated with stature. The 14 SNPs are in the vicinity of *PLAG1* and the Coiled-coil-helix-coiled-coil-helix domain containing 7 (*CHCHD7*) gene, which are 540 bp apart (Fig. 6a). The 14 candidate SNPs are shown in Fig. 6a with coordinate locations relative to HPRS-predicted regulatory regions. The HPRS database suggests a strategy for further filtering these fine-mapped SNPs in two ways, first to prioritize gene targets and second to prioritize SNPs. The design of the validation experiment by Karim et al [40] did not separate the two SNPs (rs209821678 and rs210030313) in the promoter region because both the long and short fragments used for activity assays in the study contained both SNPs. The HPRS prediction separates the two SNPs into two core CAGE peaks (Fig. 6b). The two peaks suggest two potentially separate binding sites of the transcriptional machinery. HPRS resolves the shared 540 bp promoter region into separate core promoter regions and suggests a new validation design, in which three short, directional fragments focusing more specifically on core CAGE regions (two near *PLAG1* and one near *CHCHD7* gene) can be used for functional assays of SNP genotype. Measuring promoter activity of these three constructs by using the similar promoter luciferase assay and transcription factor binding assay employed by Karim et al [40] may confirm which of the two SNPs is causative and which gene is affected.

**Figure 6.**
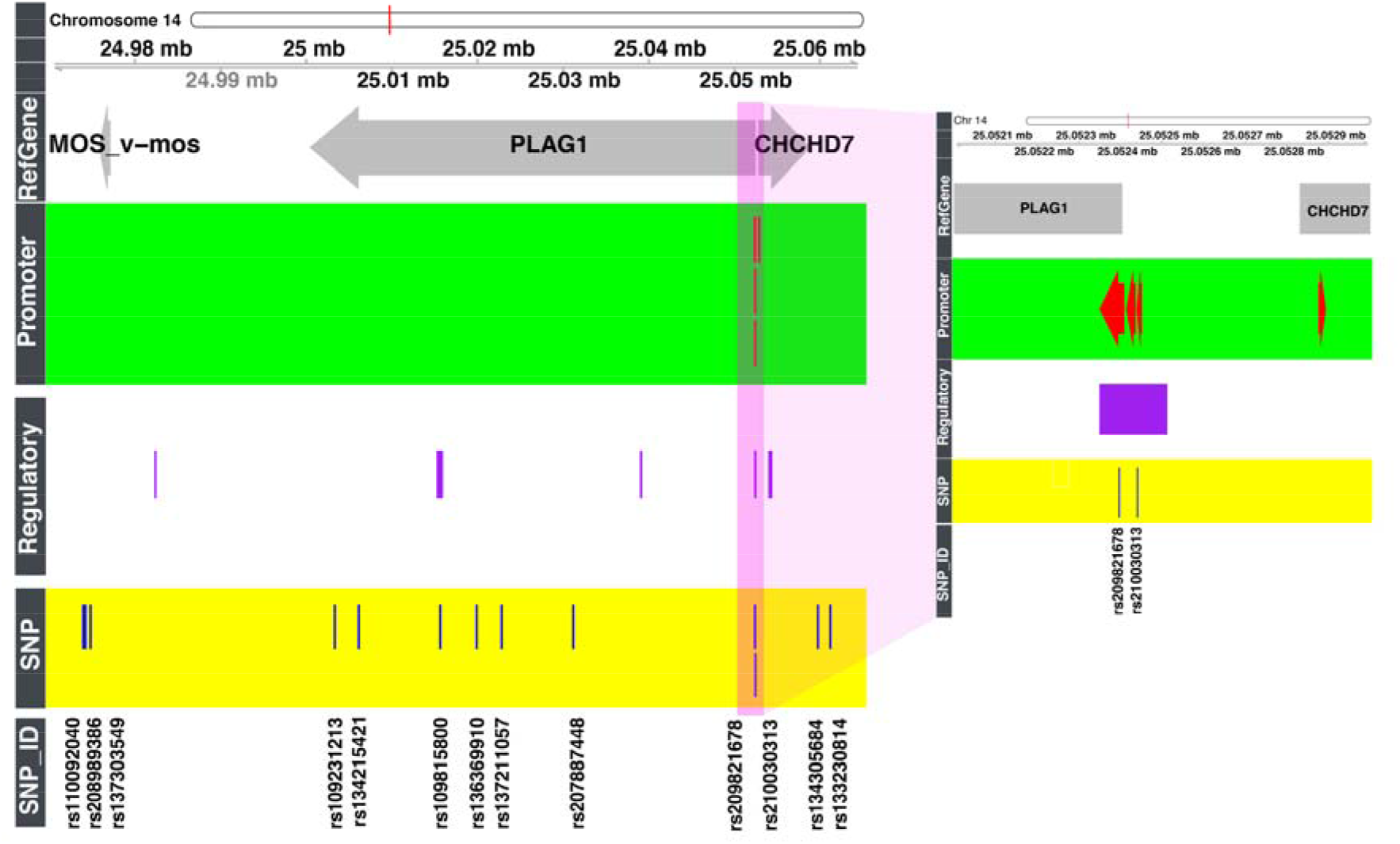
Application of the regulatory database to prioritize significant bovine SNPs identified by GWAS studies for functional validation. Overview of 13 significant SNPs fine-mapped by Karim et al [40] is shown in the left panel. Among those SNPs, only three overlap regulatory regions and promoter regions in the predicted database. A detailed view (right panel) of the two SNPs validated as causative in Karim et al [40]. Both SNPs are within promoter regions of the *PLAG1* gene, but not the *CHCHD7* gene. The Regulatory (enhancers, promoters, and transcription factor binding sites) and Promoter (only promoters) tracks display HPRS predicted regions.

Furthermore, by applying a scoring model for regulatory variants, we generated deltaSVM score for each of 97 million known bovine SNPs (see Supplementary Materials and Methods). The SNP rs209821678 had a deltaSVM score of -5.99. The score was beyond the 95^th^ percentile range of SVM scores for 97 million SNPs, suggesting that it may play an important regulatory role. Notably, the rs209821678 deletion of the (CCG)x11 to (CCG)x9 trinucleotide repeats lies in a predicted G-quadruplex and may cause changes in its structure, an event that could alter transcriptional activity [42]. In contrast, the SNP rs210030313 and rs109815800 did not have significant deltaSVM scores (0.51 and 3.2, respectively).

We then asked if the regions containing the SNPs interact with additional genes distant from the *PLAG1* locus. We applied HPRS for mapping interactions defined by chromatin conformation capture data (5C and Hi-C in the ENCODE human datasets) to predict distal targets of the promoter regions in the PLAG1 locus [43, 44], we found that rs209821678 and rs210030313 are within the anchor A_447043 (chr14:25,044,319-25,054,287, UMD3.1) with a predicted target region (chr14:25,478,861-25,497,096) near the *IMPAD1* (Inositol Monophosphatase Domain Containing 1). Variants within IMPAD1 have been implicated in short stature and chondrodysplasia (Table S2). Interestingly, the leading SNP identified in an analysis of pleiotropic genes affecting carcase traits in Nellore cattle, rs136543212 at chr14: 25,502,915, is slightly closer to *IMPAD1* [45]. The rs109815800 SNP, on the other hand, does not lie in any mapped Hi-C region. Together, the HPRS predicted results strongly suggest that the rs209821678 variant is the causative SNP among the 14 candidates fine-mapped by Karim et al [40].

Another example of applying the HPRS databases for analysis of non-coding mutations is for the case of the “Celtic mutation”, which causes the polled phenotype. The mutation is a 202-bp-indel, where the duplication of a 212 bp region (chr1:1705834-1706045) replaces the 10 bp (chr1:1706051-1706060)[46, 47] [48] (**Fig. 7**). The mechanism for the Celtic mutation is unknown, although it may affect the expression of *OLIGO1, OLIGO2, CH1H21orf62* and two long non-coding RNAs (*lincRNA1* and *lincRNA2*) [46, 47]. We found that the whole 10 base deletion, but not the upstream 212 base duplication, is within an HPRS predicted enhancer sequence (chr1:1706046-1706182, UMD3.1). A detailed transcription factor binding motif analysis of the polled mutation site suggests that a binding site for the TF HAND1 (Heart And Neural Crest Derivatives Expressed 1) is lost due to the 10 bp deletion in animals containing the Celtic mutation (Fig. 7c). The neural crest cells give rise to the craniofacial cartilage and bone [49], suggesting that the loss of the HAND1 putative binding site is a plausible explanation for the altered craniofacial development in Polled animals. Additionally, using information from Hi-C in the human genome [44], we found the mutation is within a mapped interaction targets of the regions Hi-C A_264635 (chr1:1706078-1714122, UMD3.1) and A_264636 (chr1:1698252-1706077, UMD3.1) and interacts with genes 100s of Kb away (Fig. 7: bottom panel, and Table S2). Although, the above hypothesis requires experimental validation, it shows that applying HPRS approach could lead to biological hypothesis for underlying effects of causative mutations within noncoding regions.

**Fig. 7.**
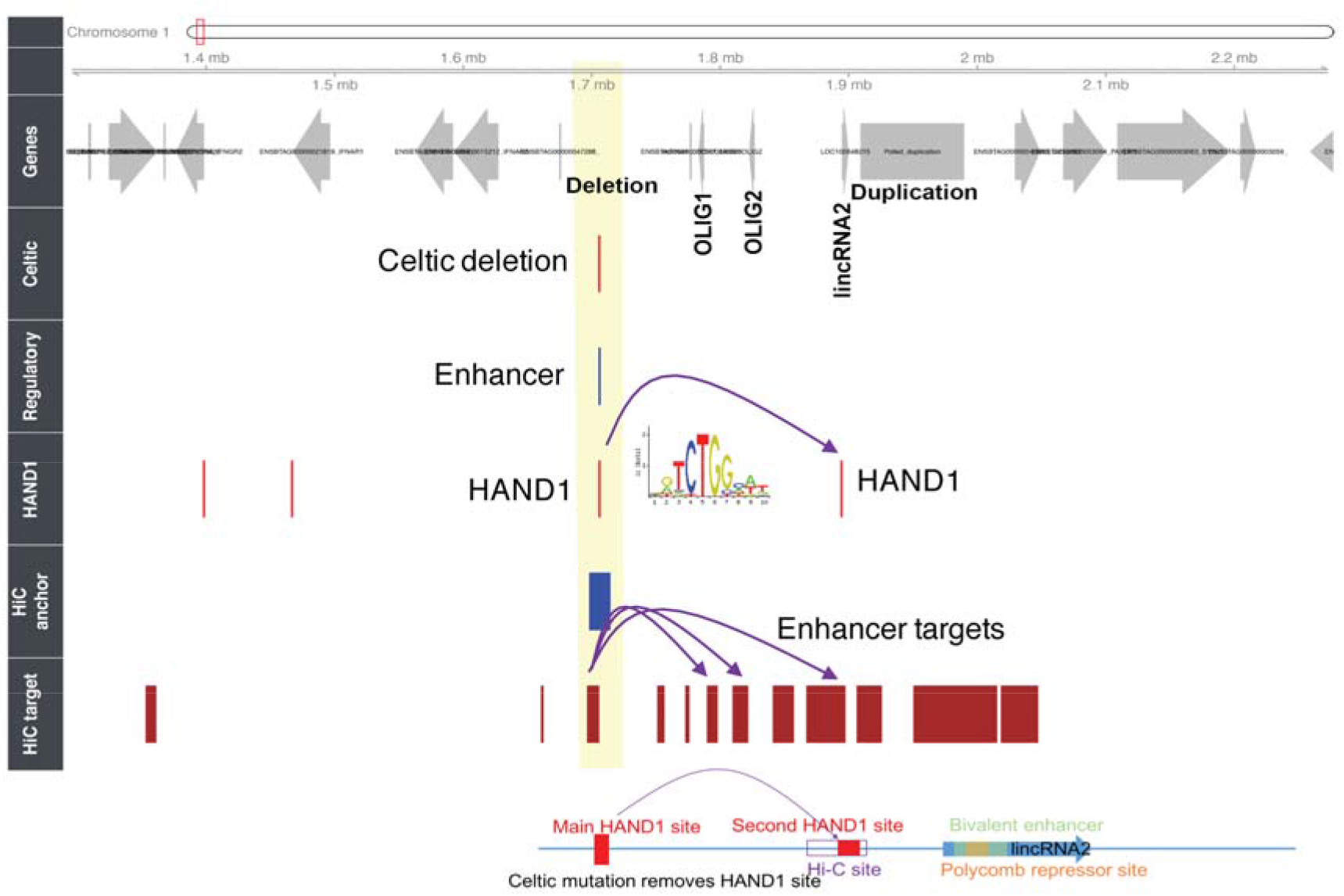
A potential model for effect of the Celtic mutation. Using human Hi-C (chromosome conformation capture) data and scanning of transcription factor binding sites, we generated a hypothesis to predict cattle regulatory targets for polled mutation (HiC target, HiC anchor and HAND1 tracks). The Regulatory (enhancers, promoters, and transcription factor binding sites) tracks display the HPRS predicted region. Two common mutations on chromosome 1 in cattle have been associated with polled cattle. One is a 202-bp-indel (“Celtic mutation”). The other is an 80 kb duplication ~300 kb away. Purple arrows on the top link the Hi-C anchor to multiple targets mapped from human to cattle genome. Map with exact size and location of the regulatory regions and the Hi-C anchor overlapping the Celtic mutation and its targets.

Therefore, from the two examples described above (and from the Callipyge example described in the supplementary section), we found that the HPRS regulatory database can be used to prioritize SNPs and genetic variants that were identified by GWAS studies and to draw hypotheses about biological mechanisms of a causative SNP.

### Limitations of the methods

The main aim of the HPRS pipeline is to predict as many regulatory regions and as accurately as possible, so that the dataset could be applied for functional SNP analysis in the target species. However, given the uncertain nature of promoter and enhancer identification, the rate of false positives and negatives by HPRS is difficult to determine. In our analysis, all of the reference cattle liver enhancers were included in the initial unfiltered datasets, although ~25% were lost during the filtering process. Similarly, 96% of reference cattle liver dataset promoters were covered by the unfiltered dataset, with less than 3% lost in the filtering process. A limitation of the HPRS filtering process is the requirement to use a species-specific data set. Nevertheless, compared to the large number of datasets and biochemical assay types that are required to create a comprehensive coverage of regulatory regions, the number of species-specific datasets needed for HPRS is small. In this paper, for each of the three species (mouse, cattle and pig), we used data from only three biological replicates of H3K27Ac assay, which was generated within a scope of one project as reported in Villar et al [22], to successfully filter the Universal Dataset. In addition, the approach cannot predict promoters and enhancers that are unique to the species, for example promoters and enhancers that are present in the cow, but not present in humans. These unique promoters/enhancers are likely to be a small proportion of the total promoter/enhancer set. Indeed, the lineage specific promoters and enhancers across 20 mammalian species were around 5 % of the total promoters and enhancers [22]. Of note, relevant human input datasets can be integrated depending on the aim of an analysis. For example, if the focus is to study milk production, the HPRS pipeline can be applied for more relevant tissues, such as the mammary gland. Future cattle-specific datasets can be incorporated into the HPRS pipeline to address the tissue and species specificity issues.

In contrast to the HPRS pipeline prediction of regulatory regions, the prediction of causative genetic variation within regulatory regions is much more challenging. The current approach relies on the enrichment of sequence motifs within regulatory regions relative to non-regulatory regions. At least some of the motifs are TFBSs, but there are likely to be other types of motifs, such as G-quadraplexes, present in regulatory regions. While the predicted datasets can be useful for generating relevant hypotheses, the identification of causal variants still requires considerable future refinement and validation.

## Conclusions

We have developed the HPRS pipeline using a large collection of existing human genomics data and a limited number of cattle specific datasets to predict a database of cattle regulatory regions that covers a large number of active promoters, enhancers and TFBSs. The database generated here is not a final product because HPRS is capable of readily integrating new cattle-specific datasets into its mapping and filtering pipeline to expand, refine and validate the databases. Moreover, the HPRS pipeline can be applied to data of other mammalian species and by scientists without computer programming skills. We anticipate that the pipeline will be used to integrate large-scale datasets from the FAANG consortium, when they become available, with complementary data from human research. The immediate application of the regulatory database is to complement the current species specific GWAS analysis by (1) discovery of potential regulatory mechanisms of SNPs lying outside gene coding regions, (2) prioritising SNPs that are statistically significant at a genome-wide level but located within regulatory regions, (3) prioritising SNPs that are at low allele frequency but have potential for large effects, and (4) suggesting possible causative SNPs as targets for precise genome editing or selective breeding practices.

## Methods

The complete HPRS pipeline is divided into three modules: mapping, filtering, and SNP analysis. The whole pipeline and documentation are available from the CSIRO BitBucket [50].

### HPRS mapping pipeline

We developed a mapping strategy based on four elements: (1) selecting a suitable combination of human databases as HPRS inputs; (2) finding an optimal sequence identity threshold in the target genome; (3) finding options to remove less confident mapped results, and; (4) adding multiple mapped regions that meet a high sequence similarity threshold. Depending on the species, targeted tissues or regulatory categories of interest, users can select suitable human databases using the following suggested criteria: types of regulatory regions (promoters, enhancers, and TFBSs), biochemical assays, computational models for combining data, and data sources (tissues, cell lines, traits). Second, by applying the UCSC liftOver tool [23], regions that were aligned at genome-scale (by LastZ pair-wise genome alignment [51]) were fine-mapped to identify target regions with proportion of sequence identity to the original regions (minMatch) higher than a selected cut-off. We recommend an optimal minMatch=0.20 and not allowing multiple mapping for this step. Users can vary input parameters (minMatchMain and minMatchMulti) in the HPRS mapping script (Main_Mapping_Pipeline.py) to optimize the minMatch suitable to specific datasets that may have different features such as sequence length and conservation. Third, mapped regions resulting from using a low minMatch cut-off (0.20) were filtered to retain only regions with exact reciprocal mapping back to human genome, with the condition that both the left and right borders of the reciprocally mapped regions were within 25 bp windows of the original regions. Fourth, to accommodate regions possibly resulting from duplication events, the HPRS mapping pipeline added a step to remap regions that are unmapped or are not reciprocally mapped by allowing multiple mapped results to be included while setting a high sequence similarity threshold (specified by the minMatchMulti parameter, ≥ 0.80). Fig. S1a shows some of the expected mapping scenarios.

In addition to the customized minMatchMain and minMatchMulti parameter inputs, the Main_Mapping_Pipeline.py script also takes user-specified chain files for target species, which can be any of the mammalian species with chain files available from the UCSC databases or generated in-house. The HPRS mapping pipeline enables fast mapping of as many databases as necessary. The script PostHPRSMapping_MergeDifferentDatabaseTypes.py (available in the CSIRO BitBucket [50]) can be used to combine resulting datasets into one dataset containing non-overlapping regions. For example, we merged enhancer databases from 88 ROADMAP tissues/primary cell lines, and five additional promoter, enhancer and TFBS databases. The script also collapses names of overlapping regions into a comma separated field that can be used to count the total number of annotations for each merged region.

### HPRS filtering pipeline

Detailed description of the seven filters is presented in the Supplementary Materials and Methods section. Briefly, the HPRS filtering pipeline was written in R and contains seven filtering steps (Fig. 4, Table 4). The input file is a merged metadata file, in which each region was calculated for the number of CAGE peaks mapped, the RNA-Seq signal from 86 cattle RNA-Seq datasets, the Villar H3K27Ac signal, the SVM enhancer scores (enhancer activity predicted by a machine learning classification method, gkmSVM) [52], the number of overlapping annotations, the conservation score based on the UCSC 100 way vertebrate alignment [53], and the number of TFBSs based on Cluster-Buster scanning [54]. The main filtering pipeline was HPRS_Filtering_pipeline.Rmd. We tested a range of parameters and recommend using the parameters set in the script. In addition, prior to running this main script, users can choose to optimize parameters suitable to specific datasets using the script HPRS_Filtering_optimize_FilterOrder.Rmd, which calculates RatioP and RatioE (average number of enhancers and promoters per Mb of the total length of all predicted enhancers and promoters) for each filter and for a range of filter parameters so that the optimal parameters are used in the main filtering pipeline. The filtering pipeline was written in a way that it is simple to add or remove filter layers depending on availability of species-specific data.

### Methods to apply HPRS dataset for regulatory SNP analysis

The HPRS dataset can be applied for the selection of top candidate SNPs in regulatory regions which are present in existing genotyping SNP chips. The selected SNPs form a small set of SNPs that are more likely to be causal or associated to phenotypes. Using these SNPs for GWAS analysis may reduce noise compared with using a large number of non-causal but in high linkage disequilibrium to causal SNPs. The top candidate SNPs can be selected by the identification of SNPs belonging or not belonging to the following categories: the Universal Dataset; the Filtered Dataset; the TFBSs of the predicted regulatory regions; and regulatory regions active in tissues related to the trait of interest. In addition, deltaSVM scores can be used as one of the indicators for potential SNP effects, as discussed in the supplementary method section. Alternatively, the dataset can be used for post-GWAS analysis, in which significant SNPs in non-coding regions that are identified from GWAS can be assessed for potential effect on gene regulatory activity. We have discussed examples of applications for the cases of pleiotropic SNPs, climatic adaptation associated SNPs, and associated SNPs milk-production traits (Fig. S1, Table S1), and of post-GWAS analysis for the stature phenotype and callipyge phenotype (Fig. 6 and Tables S2, S3).

We developed an implementation pipeline of the gkm-SVM model to estimate SNP effects on enhancer activities in cattle by adapting the model to the case where very limited species-specific ChIP-Seq data are available for model training (See Supplementary Materials and Methods).

### Data availability

We have made all HPRS Python, R and Shell scripts publically available with usage instruction from the CSIRO BitBucket [50]. These codes can be used to perform all steps from mapping, to filtering and scoring regulatory SNPs. Supporting data and scripts to run the HPRS pipeline are also available via the *GigaScience* database, GigaDB [51].

All human databases used for prediction are publically available (Table S5). Results of predicted regulatory regions, including the Universal Datasets and the filtered datasets, for cattle and pig are available as Supplementary Materials of this article. For cattle, we provide deltaSVM scores for ~97 million SNPs, which can be used as one of the parameters for assessing potential SNP effects. Additionally, we share predicted Universal Datasets (not yet filtered) for ten other mammalian species in a format compatible for uploading to the UCSC genome browser (Table 5 and Fig. S7). These 10 additional datasets can be useful for exploring potential regulatory effects from non-coding genomic regions.

## Acknowledgm ents

We thank Dr Tony Vuocolo (CSIRO) for insightful discussions, Dr Li Congjun (ARS, USDA) for sharing the update of a published ChIP-Seq dataset, and Dr Derek Bickhart (ARS, USDA) for providing us with the updated TFBS dataset. We thank the generators of the human genomics, transcriptomics and epigenomics resources, which are highly valuable to the broader research community.

## Authors’ contributions

BD, RT, JK, BB, and QN conceived the project. QN and BD designed the HPRS algorithm. QN wrote the pipeline. QN, MNS, LPN, AR, and BH contributed to the analysis. QN and BD wrote the manuscript, with inputs from all other co-authors.

